# Divergent excitatory and inhibitory signaling in a head direction circuit

**DOI:** 10.64898/2026.01.18.700161

**Authors:** Jorin Eddy, Ali H Shenasa, Paulette Monroy Alfaro, Maria Fernanda Viveros, Daniel B Turner-Evans

**Affiliations:** University of California, Santa Cruz, Department of Molecular, Cell and Developmental Biology

**Keywords:** head direction, phase-based computing, glutamate signaling

## Abstract

Neural cell types are often clustered into inhibitory, excitatory, or neuromodulatory populations. However, the signaling mechanism between two neurons ultimately depends on their neurotransmitter-receptor pairings. Here, we demonstrate an example of a glutamatergic population of neurons that appears to either excite or inhibit their downstream partners depending on the type of glutamate receptor expressed in the downstream cell type. The upstream population encodes a sinusoidal head direction signal. The downstream partners that are excited by glutamate encode the same signal while the downstream partners that are inhibited by glutamate encode a 180° phase-shifted copy. The upstream population therefore appears to pass on two different phases of the same signal by making either excitatory or inhibitory connections to different downstream partners.

## Introduction

Dale’s Law states that a given neuron releases the same neurotransmitter(s) at each of its synapses. Theoreticians often use Dale’s Law to label neurons as purely excitatory or purely inhibitory. However, single neurons can simultaneously make both excitatory and inhibitory connections dependent on the composition of receptors at postsynaptic sites. Canonical examples of this phenomenon include bipolar cells in the retina and dopamine signaling pathways in the basal ganglia. Different bipolar cells have different types of glutamate receptors and thus respond differently to glutamate signaling from common upstream partners(1, 2). OFF bipolar cells depolarize in response to light-off via excitatory ionotropic glutamate channels(3), while ON bipolar cells are hyperpolarized via metabotropic glutamate channels, leading to a light-on response to the same glutamate signal. In the basal ganglia, medium spiny neurons (MSN) in the direct pathway predominantly express D1 dopamine receptors, which increase the likelihood of their firing when activated, while MSNs in the indirect pathway express D2 dopamine receptors, which decrease the likelihood of firing. Differentially regulating these two pathways allows dopamine to modulate an animal’s movements(4, 5). Neurons in the mammalian brain can also release more than one neurotransmitter(6), which could lead to different downstream responses dependent on receptor composition. Thus, a single neuron could satisfy Dale’s law while simultaneously driving either excitatory or inhibitory responses in different downstream partners with different receptor compositions. Such heterogeneous signaling could play a key role in neural computations by selectively changing the “sign” of a signal as it transmitted across a synapse.

The “sign” of a population encoded signal is particularly important in phase-based computing. In phase-based computing, the phase of a sinusoidally varying signal carries information. Inverting a sinusoid produces another sinusoid that is 180° phase shifted. Different compositions of post-synaptic receptors could therefore lead one downstream partner to directly inherit the phase of its presynaptic partner (via excitation) while another would receive a 180° phase shifted version of that signal (via inhibition, i.e. by inverting the sign).

Phase-based computing is ubiquitous in neural circuits for navigation. Head direction cells use phase to encode the animal’s head direction in their population activity(7), and phase sets the properties of grid cells and vector object cells(8, 9). Phase-based signaling is also a prominent feature in the *Drosophila melanogaster* navigation system, where the phase of activity across different populations of neurons encodes the animal’s egocentric or allocentric traveling directions (Figure 1 A-C)(10–12). This heading signal originates in the EPG neurons, where it is encoded in a localized bump of activity (Figure 1 B)(13). The EPG neurons then synapse onto the glutamatergic Δ7 neurons (Figure 1 D,E)(12, 14), which convert the activity into a cosine-shaped signal before passing it to downstream partners(10, 12).

**Fig. 1.**
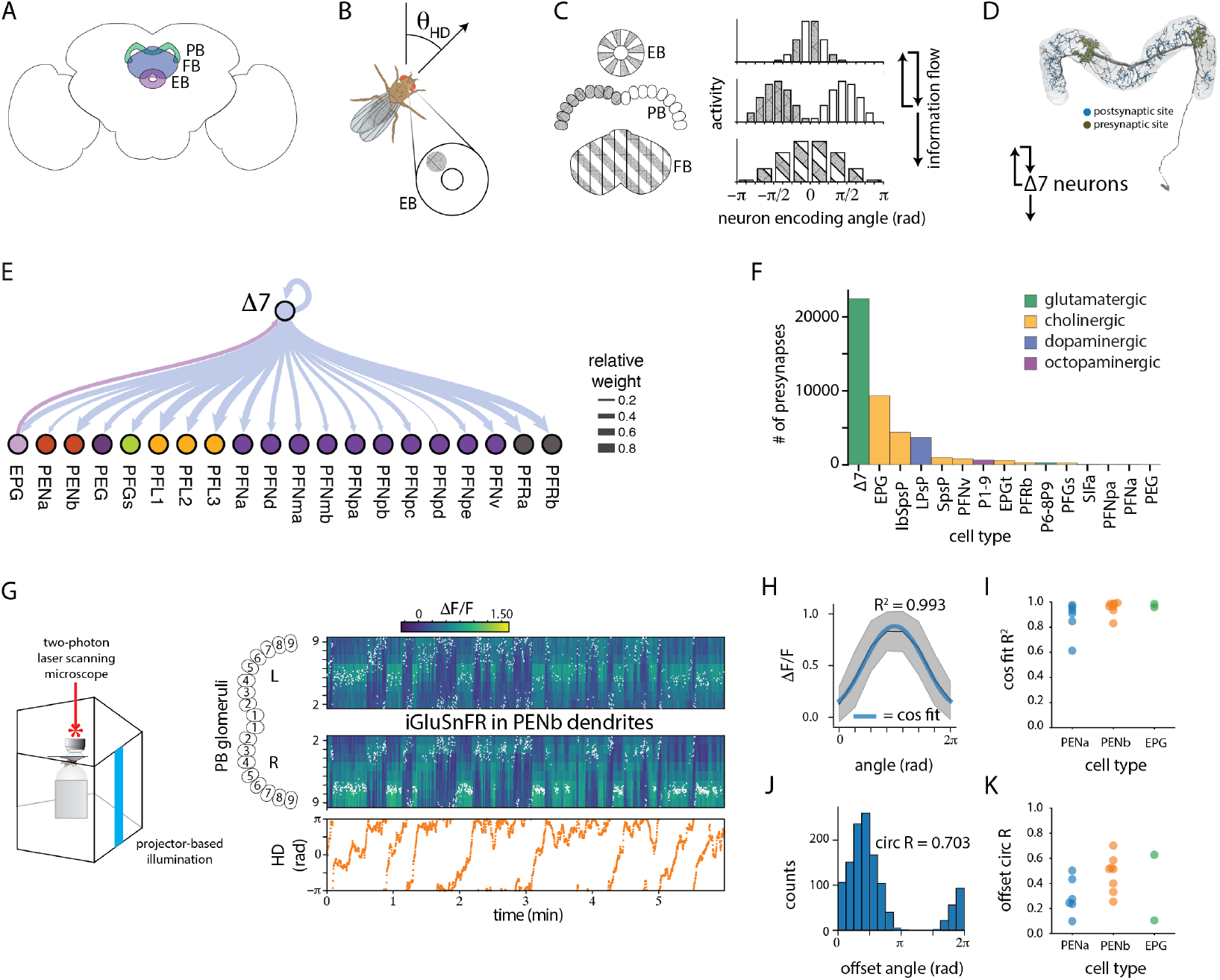
The Δ7 neurons format the Drosophila head direction signal into a cosine-shaped glutamate release profile. **A**. The Drosophila navigation center (the central complex) includes the protocerebral bridge (PB), fan shaped body (FB), and ellipsoid body (EB). **B**. Neurons in the EB encode the fly’s head direction angle (*θ*_*HD*_) in a bolus of activity that moves around the ring-shaped structure as the animal turns. **C**. The head direction signal passes between the EB and the PB and is sent from the PB out to the FB. **D**. The Δ7 neurons arborize in the PB. The density of their postsynaptic sites follows a cosine-shaped pattern across the PB while their presynaptic sites are spatially restricted to two or three discrete glomeruli. They receive input from the EPG neurons, which project from the EB to the PB, and they output to both EB and FB projecting neurons. **E**. The relative weight of connections between the Δ7 neurons and other neurons that arborize in the PB. A relative weight of 1 implies that all presynaptic connections in the PB come from the given partner. (12). **F**. The number of presynaptic sites in the PB by cell type. Each column associated with a given cell type is color coded by the known or predicted neurotransmitter that it releases. (14). **G**. Glutamate levels in the PB were monitored by expressing iGluSnFR in the dendrites of neurons postsynaptic to the Δ7 neurons. (left) Fluorescent levels were measured in head fixed flies walking on an air suspended ball while being shown a vertical stripe. The stripe’s angular position is coupled to the ball in closed loop. (right, top) iGluSnFR fluoresence in the right and left halves of the PB. The white dots show the circular mean at each time point. (right, bottom) The fly’s head direction angle over time. **H**. The mean iGluSnFR fluorescence profile across the PB for the trial shown in G. The grey shaded area shows the standard deviation while the blue curve is a cosine fit to the profile. The R2 of the fit is 0.993. **I**. The mean R2 for a cos fit to the mean iGluSnFR activity across flies. iGluSnFR was expressed in one of three different cell types that receive input from the Δ7 neurons in the PB. **J**. A histogram of the angular offsets between the circular mean of the activity (averaged across the right and left halves of the PB) for the example shown in G. The mean resultant vector length (circ R) for this distribution is 0.703. **K**. The mean circ R for the offset distributions across flies.

The unique morphology of the Δ7 neurons is believed to be responsible for this cosine conversion(12, 14). The Δ7 neurons arborize throughout the 18 discrete glomeruli of the protocerebral bridge, with broad dendritic projections and axons that selectively output in 2 glomeruli (Figure 1 D). These 2 glomeruli are separated by 7 glomeruli (hence the cell type’s name)(15). The Δ7 dendrites, on the other hand, get graded, distribution input from EPG neurons in multiple, adjacent glomeruli. This broad, graded input and selective, local output allows the population of Δ7 neurons to transform a localized bump of input activity into a sinusoidal pattern of output activity.

However, while the downstream partners of the Δ7 neurons exhibit sinusoidal population activity patterns, the phase of this sinusoid matches the Δ7 activity in some partners while others exhibit a 180° phase shifted copy of the signal(14, 16). We therefore hypothesized that different compositions of glutamate receptors in the downstream partners leads some of them to directly receive the signal via excitation while others receive the 180° phase shifted signal via inhibition. We tested this hypothesis by characterizing the profile of glutamate release across the dendrites of the downstream partners with a glutamate-sensitive fluorescent reporter, measuring the partners’ calcium activity in response to local release of glutamate, and localizing different types of ionotropic glutamate receptors on the downstream dendrites via genetic and molecular techniques. Consistent with our hypothesis, we observed that some of the downstream partners are excited by glutamate while others are inhibited and that the downstream neurons’ glutamatergic receptor composition varies in accordance with these differential responses. Taken together, our data reveal a single class of neurons that appears to be either excitatory or inhibitory dependent on the downstream receptor composition.

## Results

### The Δ7 neurons are the primary glutamatergic cell type in the protocerebral bridge, and downstream partners of the Δ7 neurons receive a cosine-shaped glutamate signal on their dendrites

The protocerebral bridge (PB) acts as a relay center that passes the head direction signal from the EPG neurons to downstream partners through the Δ7 neurons(12). The Δ7 neurons receive input from the EPG neurons and output to 25 different downstream partners (Figure 1 E). They are the primary source of input for neurons with dendrites in the PB, accounting for 51% of all labeled presynaptic sites and 99% of all glutamatergic synapses (Figure 1 F) in the hemibrain dataset(17).

As the Δ7 neurons are the primary source of glutamate in the PB and as their connectivity is graded in a way that suggests they format the EPG head direction activity into a cosine-shaped activity pattern(10, 12, 14), we predicted that the glutamate profile in the PB would also form a cosine. We therefore expressed iGluSNFr, a glutamate-sensitive fluorescent reporter, in the dendrites of three cell types that receive input from the Δ7 neurons in the PB: the EPG, PENa, and PENb neurons. We then monitored the glutamate signal in these neurons’ dendrites in head-fixed, walking flies with closed loop control over a vertical stripe (Figure 1 G). In each of these cell types, the iGluSNFR signal was indeed well fit by a cosine (Figure 1 H,I), and the phase of the cosine tracked the animal’s head direction at a fixed offset that varied across flies (Figure 1 J,K).

### Some downstream partners of the Δ7 neurons are inhibited by glutamate while others are excited

The downstream partners of the Δ7 neurons can have different activity profiles in the PB, with phases that are 180° shifted from one another. Specifically, the PENb neurons were previously shown to have an activity profile that is 180° out of phase with the activity profile of the EPG, PENa, and PEG neurons in the PB (Figure 2 A) (14, 16). We therefore hypothesized that these different cell types would respond differently to glutamate, i.e. that the PENb neurons would be excited by glutamate while the EPG and PENa neurons would be inhibited (Figure 2 B,C). To test this hypothesis, we expressed GCaMP6s, a genetically encoded calcium indicator, in each of these neurons in turn and monitored their responses in *ex vivo* brains to local puffs of glutamate, introduced through a pulled glass capillary (Figure 2 D, see Figure 4 A for puffing conditions, which were chosen by comparing the physiological glutamate levels in walking flies to the glutamate levels introduced by puffing, as monitored by iGluSnFr, Figure 4 B). Consistent with our predictions, fluorescence in the EPG neurons decreased (Figure 2 E,F) in response to glutamate while fluorescence in the PENb neurons increased (Figure 2 G). These responses were consistent across preparations (Figure 2 H,I). Unexpectedly, the signal in the PENa neurons also increased, though not as consistently or as strongly as the signal in the PENb neurons (Figure 4 D,E). We did not measure the response in PEG neurons. To control for network effects, we washed in mecamylamine, an antagonist of nicotinic acetylcholine receptors and observed similar effects (Figure 4 D).

**Fig. 2.**
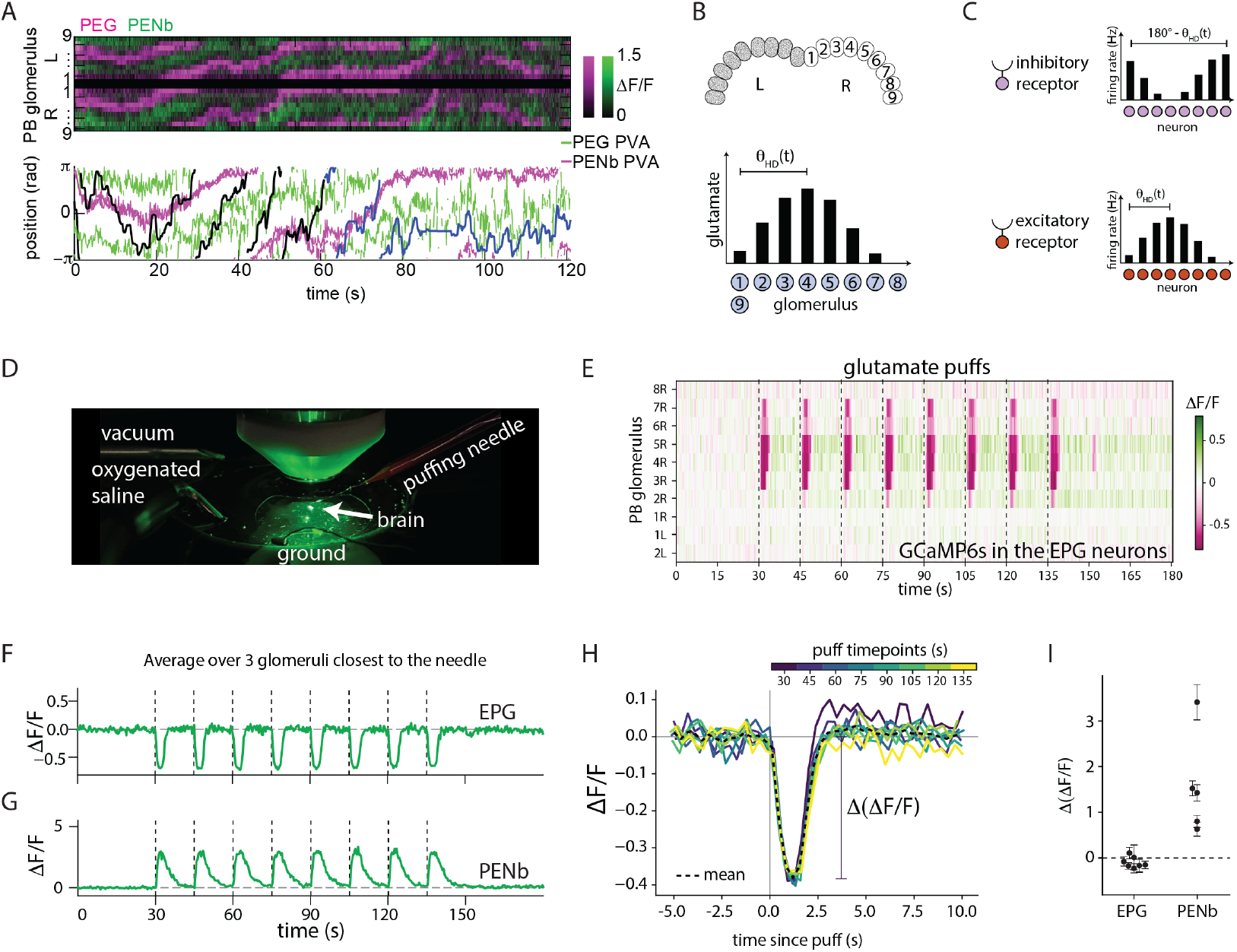
Some of the Δ7 neurons downstream partners are excited by glutamate while others are inhibited. **A**. (top) Two color imaging of the calcium activity in the PEG and PENb neurons. (bottom) The population vector average of the PEG and PENb activity overlaid on the fly’s heading. The fly is in the dark for the first 60 s (black curve) and then shown a closed-loop stripe for the remainder of the trial (blue curve). **B**. (top) The PB is divided into 18 glomeuli, 9 on the left and 9 on the right. (bottom) The head direction signal from the EB is passed to each of the 9 glomeruli. **C**. A cosine-shaped pattern of glutamate release could be converted to a cosine-shaped pattern of firing rates in columnar downstream neurons with a 180° phase shift via inhibitory receptors (top) or with the same phase via excitatory glutamate receptors (bottom). **D**. An overview of the setup used to deliver puffs of glutamate to the PB in an *ex vivo* preparation. **E**. The ΔF/F response to puffs of glutamate in glomeruli across the PB when GCaM6s was expressed in the EPG neurons. A 25 ms pulse was released at the time points noted by the vertical dotted lines. **F**. The average ΔF/F response in the three glomeruli closest to the needle for GCaMP6s in the EPG neurons for the preparation should in E. **G**. As in F for a preparation with GCaMP6s expressed in the PENb neurons. **H**. The traces in F aligned to the puff onset. The mean profile is shown with the dotted black line. The change in ΔF/F (Δ(ΔF/F)) is shown with the vertical line. **I**. The Δ(ΔF/F)) across flies. Note that the change in fluorescence is much greater for the PENb neurons (as seen in G) than for the EPG neurons (as seen in F, H).

### Different glutamate responses are correlated with different compositions of glutamate receptors

Different compositions of glutamate receptors in the downstream partners of the Δ7 neurons could lead them to exhibit different responses to glutamate release from the Δ7 neurons. We therefore asked whether the EPG neurons and Δ7 neurons have different concentrations of the inhibitory glutamate receptor, GluCl, in their dendrites (Figure 3) by using stochastic heat shock labeling to simultaneously express a membrane tag and an epitope-tag for the GluCl alpha subunit in single EPG and PENb neurons (18). Indeed, GluCl alpha subunits were strongly labeled in the EPG neurons while the PENb neurons exhibited minimal GluCl signal.

**Fig. 3.**
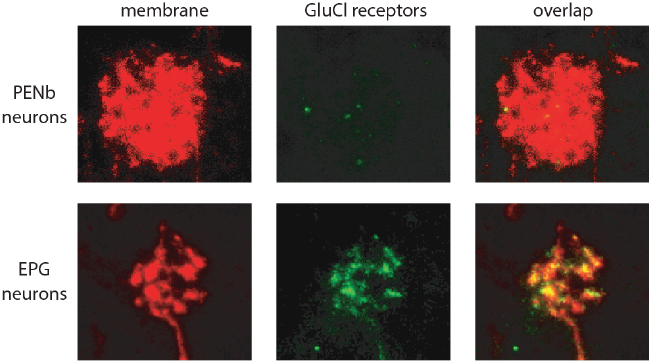
The EPG neurons, but not the PENb neurons, express tagged GluCl receptors. Labeled PB glomeruli in transgenic fly lines that express both a membrane tag (left) and an epitope-tag for GluCl receptors (middle) under a recombinase cascade in either the PENb (top) or EPG (bottom) neurons. The overlap of the membrane tag and the receptor tag is shown on the right. 7 EPG neurons across 2 brains showed strong signal while 6 PENb neurons over 2 brains showed weak signal.

**Fig. 4.**
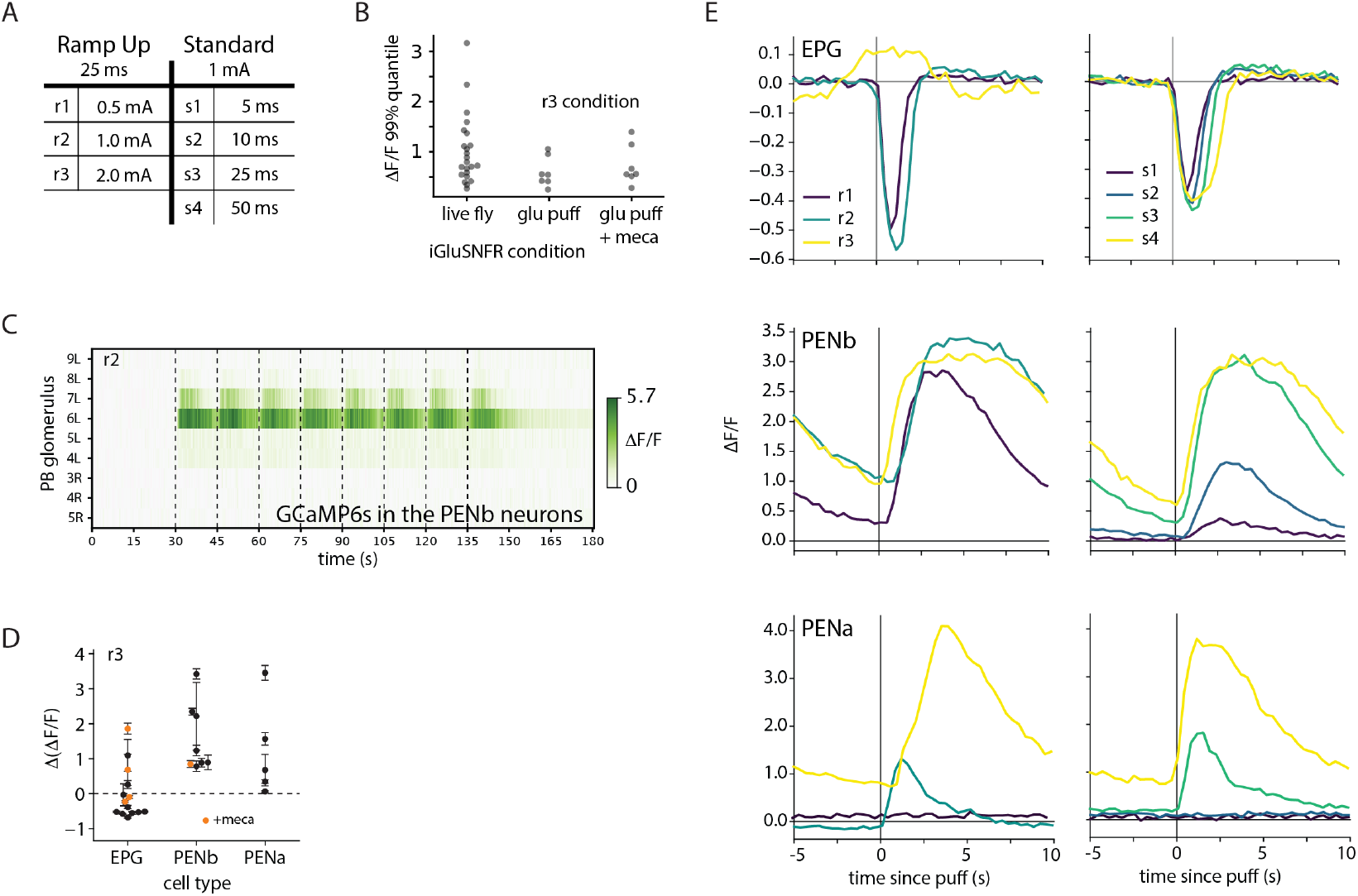
Fluorescence responses to glutamate puffing. **A**. The different conditions used to puff glutamate on the PB. **B**. The 99% quantile ΔF/F response across the PB for iGluSnFr expressed in the EPG, PENa, or PENb neurons in a live fly (left) or in an ex plant brain without (center) or with (right) mecamylamine. Responses were similar regardless of cell type, so the data points were grouped for a given condition. **C**. The ΔF/F response to puffs of glutamate in glomeruli across the PB when GCaM6s was expressed in the PENb neurons. A 25 ms pulse was released at the time points noted by the vertical dotted lines. **D**. The Δ(ΔF/F)) response across brains as shown in Figure 2 I. The x-axis denotes the neuron type expressing GCaMP6s. Both the EPG and PENb conditions also include responses in brains bathed in 50 µM mecamylamine, shown with the orange data points. **E**. Mean aligned responses across puffing conditions for 1 brain each for flies expressing GCaMP6s in the EPG neurons (top), PENb neurons (center), or PENa neurons (bottom). The ramp up (r) conditions are shown on the left, and the standard (s) conditions are shown on the right.

## Discussion

In conclusion, we observed that the primary downstream partners of the glutamatergic Δ7 neurons receive sinusoidal glutamate signals, that the EPG neurons are inhibited by local puffs of glutamate while the PENa and PENb neurons are excited, and that inhibitory GluCl alpha subunits are strongly expressed in the EPG neurons, but not in the PENb neurons.

Our results imply that one population of neurons, the Δ7 neurons, can be both excitatory and inhibitory, dependent on the composition of receptors in its downstream partners. This parallel, differential signaling mechanism is likely important for downstream computations — altering the phase of the sinusoidal signal that is passed on to downstream partners. Our results suggest that theorists should not be reticent to selectively include populations of neurons that are both excitatory or inhibitory, though we acknowledge that biological examples of this phenomena are not the norm.

We were also surprised to observe that the PENa neurons were excited by glutamate, as their activity profile in the PB was previously observed to directly follow the EPG activity (16, 19), and we would therefore expect them to be similarly inhibited. In addition to tagged GluCl receptors, we also attempted to label excitatory kainate and NMDA-like glutamate receptors in the EPG, PENa, and PENb neurons to better understand the glutamate response of each, but the results were inconclusive.

Our results also demonstrate how characterizing the biophysical properties of different cell types can reveal important computations taking place across synapses. While significant efforts have been directed towards defining the different cell types in the brain, characterizing the biophysical components of each of these cell types will be equally important to understand the properties of connections between them as well as their input-output properties.

Finally, we note that selectively knocking out subsets of receptors is a promising future direction for revealing the computational role of different neuron types. For example, the role of the PENb neurons is still unknown in the *Drosophila* head direction circuit, and the activity of these neurons has curiously been observed to be mismatched between the PB and EB. Identifying the relevant glutamatergic receptors expressed in PB in the PENb neurons, knocking them out selectively in this cell type, and observing how the head direction signal changes would provide crucial insight into the role of these neurons in the circuit.

## Methods

### Δ7 connectivity

Δ7 connectivity data was pulled from the hemibrain v1.2.1 dataset using the getTypeToTypeTable function from neuprintrExtra and plotted using the visNetwork R package with the node positions manually specified.

### PB presynaptic analysis

The number of presynaptic sites per cell type was pulled from the hemibrain v1.2.1 dataset using the fetch_neurons function from the neuprint-python package. The primary neurotransmitter for each cell type was then manually specified as follows: the Δ7 and P6-8P9 neurons as glutamatergic, the LPsP neurons as dopaminergic, and the P1-9 neurons as octopaminergic.

### Live fly preparation - iGluSnFR imaging

The live fly experiments shown in Figure 1 followed the protocols outlined in (14), with 20xUAS-SF-iGluSnFR.A184S expressed in the PENa, PENb, or EPG neurons.

### Iontrophoresis setup

A 1M glutamate solution was prepared by mixing L-Glutamic acid monosodium salt monohydrate (MSG) with Sulforhodamine 101 dye in 10 ml of ultrapure water. The glutamate solution was then passed through a 0.45 µm PTFE Luer Lock filter to ensure removal of excess glutamate salt and collected in a 15 ml conical tube.

Freshly pulled glass micropipettes were filled with 20 µl of the glutamate solution, any air bubble were removed, and the pipette was then fastened to a pipette holder. The pipette holder was then secured into a micromanipulator dock fixed to the microscope stage.

### Dissection and electrode positioning

Imaging was performed with an Olympus 2-photon microscope running Fluoview software using a 940 nm pulsed femtosecond laser. A separate laptop was connected to the ion-trophoresis pump and used to run custom LabVIEW code. This LabVIEW code controlled the puffing dynamics.

Brains were dissected in cold extracellular saline solution, and the glial sheath was removed just above the protocerebral bridge. Removal of the glial sheath was necessary for proper micropipette insertion and glutamate diffusion during the puffing trials. Brains were then mounted on a 12 mm PLL-coated coverslip and transferred together to a custom made, laser-cut microscopy dish designed to fit securely in the microscope stage.

Once secured on the stage, oxygenated saline was perfused through the dish to maintain cell activity following dissection. This was done by passing carbogen through a diffuser placed in a 50 ml conical vial containing the extracellular saline, which was drained into the dish via an IV drip line. Excess saline was removed from the dish via a vacuum positioned at the surface of the solution.

An iontophoresis pump was used to drive glutamate out of the micropipette. The ground electrode was submerged within the saline adjacent to the brain, while the wire electrode was encapsulated within the micropipette and glutamate solution. Prior to injection, −2.0 µm of current was briefly passed through the pipette into the dish to confirm that the electrophoresis pump was functioning correctly and that there were no blockages in the pipette.

Epifluorescent light was used to locate the brain and pipette tip. Once both were focused within the same Z-plane (just above the protocerebral bridge), the light path was switched to the two-photon detector. The pipette was then inserted into the brain and positioned adjacent to one arm of the protocerebral bridge such that the tip was centered on a single glomerulus. Once the appropriate capture plane was achieved, a brief “post-insertion clear” trial was run to remove any cellular debris on the tip of the pipette (three, 50 ms pulses at −2 µA each).

### Mecamylamine

For blocking possible feedback activity into the PB that would follow from the stimulation of PB neurons with the glutamate pulses, 50 µM of mecamylamine (Sigma-Aldrich, St Louis, MO) was perfused into the saline bath.

## ACKNOWLEDGEMENTS

The authors express our deep gratitude to Kerstin Richter for invaluable help with experiments, Ben Abrams for microscopy training and support, Vivek Jayaraman and the HHMI Janelia Research Campus for support in performing pilot experiments that ultimately led to this study, and Yi Zuo, Euiseok Kim, and Brad Hulse for feedback throughout the process.

## AUTHOR CONTRIBUTIONS

JE performed the glutamate puffing experiments and immunohistochemistry. AS designed and implemented the data analysis pipeline. MFV developed fly lines. PMA performed the heat shock labeling of GluCl receptors, IHC, and imaging. DBTE designed the study, provided guidance throughout, and wrote the manuscript.

## Bibliography

1. Thomas Euler, Silke Haverkamp, Timm Schubert, and Tom Baden. Retinal bipolar cells: elementary building blocks of vision. Nature Reviews Neuroscience, 15(8):507–519, 2014. ISSN 1471-003X. doi: 10.1038/nrn3783.

2. Steven H. DeVries, Wei Li, and Shannon Saszik. Parallel processing in two transmitter microenvironments at the cone photoreceptor synapse. Neuron, 50(5):735–748, 2006. ISSN 0896-6273. doi: 10.1016/j.neuron.2006.04.034.

3. C. Puller, E. Ivanova, T. Euler, S. Haverkamp, and T. Schubert. OFF bipolar cells express distinct types of dendritic glutamate receptors in the mouse retina. Neuroscience, 243: 136–148, 2013. ISSN 0306-4522. doi: 10.1016/j.neuroscience.2013.03.054.

4. Nicolas X. Tritsch and Bernardo L. Sabatini. Dopaminergic modulation of synaptic transmission in cortex and striatum. Neuron, 76(1):33–50, 2012. ISSN 0896-6273. doi: 10.1016/j.neuron.2012.09.023.

5. Charles R. Gerfen, Thomas M. Engber, Lawrence C. Mahan, Zvi Susel, Thomas N. Chase, Frederick J. Monsma Jr., and David R. Sibley. D1 and d2 dopamine receptor-regulated gene expression of striatonigral and striatopallidal neurons. Science, 250(4986):1429– 1432, 1990. ISSN 0036-8075. doi: 10.1126/science.2147780.

6. Michael L. Wallace and Bernardo L. Sabatini. Synaptic and circuit functions of multitransmitter neurons in the mammalian brain. Neuron, 2023. ISSN 0896-6273. doi: 10.1016/j.neuron.2023.06.003.

7. Brad K. Hulse and Vivek Jayaraman. Mechanisms underlying the neural computation of head direction. Annual Review of Neuroscience, 43(1):1–24, 2016. ISSN 0147-006X. doi: 10.1146/annurev-neuro-072116-031516.

8. Torkel Hafting, Marianne Fyhn, Sturla Molden, May-Britt Moser, and Edvard I. Moser. Microstructure of a spatial map in the entorhinal cortex. Nature, 436(7052):801–806, 2005. ISSN 0028-0836. doi: 10.1038/nature03721.

9. Øyvind Arne Høydal, Emilie Ranheim Skytøen, Sebastian Ola Andersson, May-Britt Moser, and Edvard I. Moser. Object-vector coding in the medial entorhinal cortex. Nature, 568 (7752):400–404, 2019. ISSN 0028-0836. doi: 10.1038/s41586-019-1077-7.

10. Cheng Lyu, L. F. Abbott, and Gaby Maimon. Building an allocentric travelling direction signal via vector computation. Nature, pages 1–6, 2021. ISSN 0028-0836. doi: 10.1038/s41586-021-04067-0.

11. Jenny Lu, Amir H. Behbahani, Lydia Hamburg, Elena A. Westeinde, Paul M. Dawson, Cheng Lyu, Gaby Maimon, Michael H. Dickinson, Shaul Druckmann, and Rachel I. Wilson. Transforming representations of movement from body-to world-centric space. Nature, pages 1–7, 2021. ISSN 0028-0836. doi: 10.1038/s41586-021-04191-x.

12. Brad K Hulse, Hannah Haberkern, Romain Franconville, Daniel B Turner-Evans, Shinya Takemura, Tanya Wolff, Marcella Noorman, Marisa Dreher, Chuntao Dan, Ruchi Parekh, Ann M Hermundstad, Gerald M Rubin, and Vivek Jayaraman. A connectome of the drosophila central complex reveals network motifs suitable for flexible navigation and context-dependent action selection. eLife, 10, 2021. doi: 10.7554/elife.66039.

13. Johannes D Seelig and Vivek Jayaraman. Neural dynamics for landmark orientation and angular path integration. Nature, 521(7551):186–91, 2015. ISSN 0028-0836. doi: 10.1038/nature14446.

14. Daniel B. Turner-Evans, Kristopher T. Jensen, Saba Ali, Tyler Paterson, Arlo Sheridan, Robert P. Ray, Tanya Wolff, J. Scott Lauritzen, Gerald M. Rubin, Davi D. Bock, and Vivek Jayaraman. The neuroanatomical ultrastructure and function of a biological ring attractor. Neuron, 2020. ISSN 0896-6273. doi: 10.1016/j.neuron.2020.08.006.

15. Tanya Wolff, Nirmala A. Iyer, and Gerald M. Rubin. Neuroarchitecture and neuroanatomy of the drosophila central complex: A GAL4-based dissection of protocerebral bridge neurons and circuits. Journal of Comparative Neurology, 523(7):997–1037, 2015. ISSN 1096-9861. doi: 10.1002/cne.23705.

16. Jonathan Green, Atsuko Adachi, Kunal K. Shah, Jonathan D. Hirokawa, Pablo S. Magani, and Gaby Maimon. A neural circuit architecture for angular integration in drosophila. Nature, 546(7656):101–106, 2017. ISSN 0028-0836. doi: 10.1038/nature22343.

17. Louis K Scheffer, C Shan Xu, Michal Januszewski, Zhiyuan Lu, Shin-ya Takemura, Kenneth J Hayworth, Gary B Huang, Kazunori Shinomiya, Jeremy Maitlin-Shepard, Stuart Berg, Jody Clements, Philip M Hubbard, William T Katz, Lowell Umayam, Ting Zhao, David Ackerman, Tim Blakely, John Bogovic, Tom Dolafi, Dagmar Kainmueller, Takashi Kawase, Khaled A Khairy, Laramie Leavitt, Peter H Li, Larry Lindsey, Nicole Neubarth, Donald J Olbris, Hideo Otsuna, Eric T Trautman, Masayoshi Ito, Alexander S Bates, Jens Goldammer, Tanya Wolff, Robert Svirskas, Philipp Schlegel, Erika Neace, Christopher J Knecht, Chelsea X Alvarado, Dennis A Bailey, Samantha Ballinger, Jolanta A Borycz, Brandon S Canino, Natasha Cheatham, Michael Cook, Marisa Dreher, Octave Duclos, Bryon Eubanks, Kelli Fairbanks, Samantha Finley, Nora Forknall, Audrey Francis, Gary Patrick Hopkins, Emily M Joyce, SungJin Kim, Nicole A. Kirk, Julie Kovalyak, Shirley Lauchie, Alanna Lohff, Charli Maldonado, Emily A Manley, Sari McLin, Caroline Mooney, Miatta Ndama, Omotara Ogundeyi, Nneoma Okeoma, Christopher Ordish, Nicholas Padilla, Christopher M Patrick, Tyler Paterson, Elliott E. Phillips, Emily M. Phillips, Neha Rampally, Caitlin Ribeiro, Madelaine K. Robertson, Jon Thomson Rymer, Sean M. Ryan, Megan Sammons, Anne K. Scott, Ashley L. Scott, Aya Shinomiya, Claire Smith, Kelsey Smith, Natalie L. Smith, Margaret A. Sobeski, Alia Suleiman, Jackie Swift, Satoko Takemura, Iris Talebi, Dorota Tarnogorska, Emily Tenshaw, Temour Tokhi, John J. Walsh, Tansy Yang, Jane Anne Horne, Feng Li, Ruchi Parekh, Patricia K Rivlin, Vivek Jayaraman, Marta Costa, Gregory SXE Jefferis, Kei Ito, Stephan Saalfeld, Reed George, Ian Meinertzhagen, Gerald M Rubin, Harald F Hess, Viren Jain, and Stephen M Plaza. A connectome and analysis of the adult drosophila central brain. eLife, 9:e57443, 2020. doi: 10.7554/elife.57443.

18. Piero Sanfilippo, Alexander J. Kim, Anuradha Bhukel, Juyoun Yoo, Pegah S. Mirshahidi, Vijaya Pandey, Harry Bevir, Ashley Yuen, Parmis S. Mirshahidi, Peiyi Guo, Hong-Sheng Li, James A. Wohlschlegel, Yoshinori Aso, and S. Lawrence Zipursky. Mapping of multiple neurotransmitter receptor subtypes and distinct protein complexes to the connectome. Neuron, 2024. ISSN 0896-6273. doi: 10.1016/j.neuron.2023.12.014.

19. Daniel Turner-Evans, Stephanie Wegener, Hervé Rouault Romain Franconville, Tanya Wolff, Johannes D Seelig, Shaul Druckmann, and Vivek Jayaraman. Angular velocity integration in a fly heading circuit. eLife, 6:e23496, 2017. doi: 10.7554/elife.23496.

